# Segregation of floricolous ants along latitudinal and urbanization gradients

**DOI:** 10.1101/588186

**Authors:** Alan Vergnes, Quentin Rome, Inès Gayral, Colin Fontaine

## Abstract

Recent call has been made to study the biogeography of species interactions in order to better understand ecosystems’ states and processes, as well as their response to global anthropogenic disturbances. Ants (Formicidae) are a dominant group of arthropods with a central role in ecosystem functioning. Many ant species, those feeding on liquids, are floricolous and consume nectar. The biogeography of ant-flower interactions is still poorly studied and especially in temperate area. Here we quantify variations in ant-flower interaction frequency in response to latitudinal and urbanization gradients at a country scale.

We used data from a flower-visitor monitoring program that includes pictures on 2511 flower plants across Continental France (Mainland) and over 4 years. We analysed the occurrence of the ant-flower interactions along two gradients: latitude and urbanization, this for 10 ant taxa corresponding to different taxonomical level (from family to species).

Ants visited 26 % of the sampled plants. Most of the observed ant-flower interaction involved the subfamily Formicinae (82.1 %), followed by Myrmicinae and Dolichoderinae (6.9 % and 4.6 % respectively). Globally, (i) the probability of occurrence was negatively related to latitudes (ii) and to urbanization at lower latitude. (iii) Responses to latitude among sub families, genus and species level responses were segregated and taxonomically aggregated.

At lower taxonomic levels we found clear latitudinal niche partitioning among ant taxa suggesting that competition, on both evolutionary and ecological time scales, is a major process structuring ant communities. Finally, our results highlight that the effects of large scale perturbation like urbanization can vary and affect latitudinal gradient.

## Introduction

A major aim of biogeography is to unravel the spatio-temporal distribution of species and ecosystems as well as the underlying abiotic factors and biotic processes shaping such distribution (Lomolino *et al.* 2006). This scientific field provided major insights for the understanding of natural gradients such as the decrease of diversity with increasing latitude at global scale (Gaston 2000, Rolland *et al.* 2014). There is growing evidence that global change can act as an additional driver shaping such large-scale diversity patterns. Indeed, human activities induce major environmental modifications such as climate or land use changes, with consequences on both species distribution and the functioning of species assemblages (Sala *et* al. 2000, Tylianakis *et al.* 2008). Understanding how such natural and human induce drivers of species distribution interact appears as one of the current challenges of biogeography (Parmesan *et al.* 2005).

Here we propose to address this challenge investigating the combined effects of latitude and urbanization on the distribution of the interaction between liquid-feeding ants and flower plants across Continental France.

Urbanization represents the prime driver of land use change in Europe (EEA 2010), with city areas increasing at a high rate and projected to double by 2030, reaching 6 % of emerged earth’s surface (Banque_Mondiale 2009). Urbanization has consequences far beyond cities’ boundaries and impacts biodiversity patterns at a large scale (Pickett *et al.* 2011) mainly causing an impoverishment of communities and increasing the dominance of few urbanophile species (McKinney 2006).

For ants, most studies showed a negative effect of urbanization on species richness (Antonov 2008, Antonova and Penev 2006, Forys and Allen 2005, Holway and Suarez 2006, Lessard and Buddle 2005, López-Moreno *et al.* 2003, Majer and Brown 1986, Sanford *et al.* 2009, Thompson and McLachlan 2007, Uno *et al.* 2010, Xu *et al.* 1998) and one study a positive effect (Ives *et al.* 2011). The effects of urbanization on ant’s abundance have been less studied and showed a more contrasted pattern (Antonova and Penev 2006, Buczkowski and Richmond 2012, Sanford *et al.* 2009, Uno *et al.* 2010). Indeed, rare species seem to exploit the modifications induced by urbanization and can reached high densities in cities (McKinney 2008, Vepsäläinen *et al.* 2008). To our knowledge, only one study has investigated the response of ant-plant interaction (Thompson and McLachlan 2007) which highlighted an increase of seeds’ removal rate by the few species that remained in more urbanized forest.

Ants (Hymenoptera, Formicidae) are among the most abundant and ecologically significant groups on earth, sometime referred then as “the little things that rule the world” (Hölldobler and Wilson 1990), as they can deeply modify the environment, to the point of being considered as ecological engineers (Jones *et al.* 1994). As suggested by the correlated pattern of diversification between ants and angiosperm (Moreau *et al.* 2006), the interactions establish by ants with flowering plants might have been key to their current widespread distribution (Rico-Gray and Oliveira 2007). Their distribution varies along latitudinal gradient (Baroni Urbani and Collingwood 1977) and so their taxonomic diversity at both global (Dunn *et al.* 2009, Jenkins *et al.* 2011) and local scales (Del Toro 2013, Segev 2010).

Ant-plant interactions are diverse and distributed along an antagonism-mutualism continuum, with some ants feeding directly or indirectly on plants, inhabiting plants, protecting plant against herbivory, dispersing seeds, and even pollinating plants (Rico-Gray and Oliveira 2007). For species referred as liquid-feeding, which are frequent flowers visitors, consumer-resource interactions seem dominant. They often directly feed on plants, either from extrafloral nectary (Rico-Gray and Oliveira 2007) or within flowers (Herrera *et al.* 1984). Another explanation for the presence of ants on flowers is the mutualistic association between some ants and aphids. Indeed, many liquid-feeding ants plants have developed trophobiotic relationships with homopterous (aphids, coccids, membracids or lycaenids) and collect honeydew, such as the workers of the subfamilies Formicinae and Dolichoderinae and a few species in the genus *Myrmica* and *Tetramorium* of Myrmicinae (Stadler and Dixon 2005). While protecting aphids from potential parasitoids and predators they often patrol on flowers.

Using a large dataset coming from a citizen science program monitoring flower visitors across Continental France, we proposed to analyze the combine effect of latitude and urbanization on the presence of ants on flowers. According to spatial scale and taxonomic hierarchy, latitudinal gradients of biodiversity can show various patterns (Gaston 1996, Willig *et al.* 2003): from positive to negative linear relationships, but also quadratic or non significant ones. In France, for example, Lobo *et al.* (2002) showed a quadratic relationship of dung beetles occurrences. We test whether ants’ response to latitude is linear or quadratic and to what extent such latitudinal response is affected by urbanization. We carry out our analysis at various taxonomic levels, from family to species, looking for similarities and singularities in these responses.

## Material and Methods

### 1. Studied area

The study took place across Mainland Continental France (here after France), spanning a latitudinal gradient from 51.08° to 42.32° N (Fig. 1). It covers an area of 675,000 km^2^.

**Fig 1.**
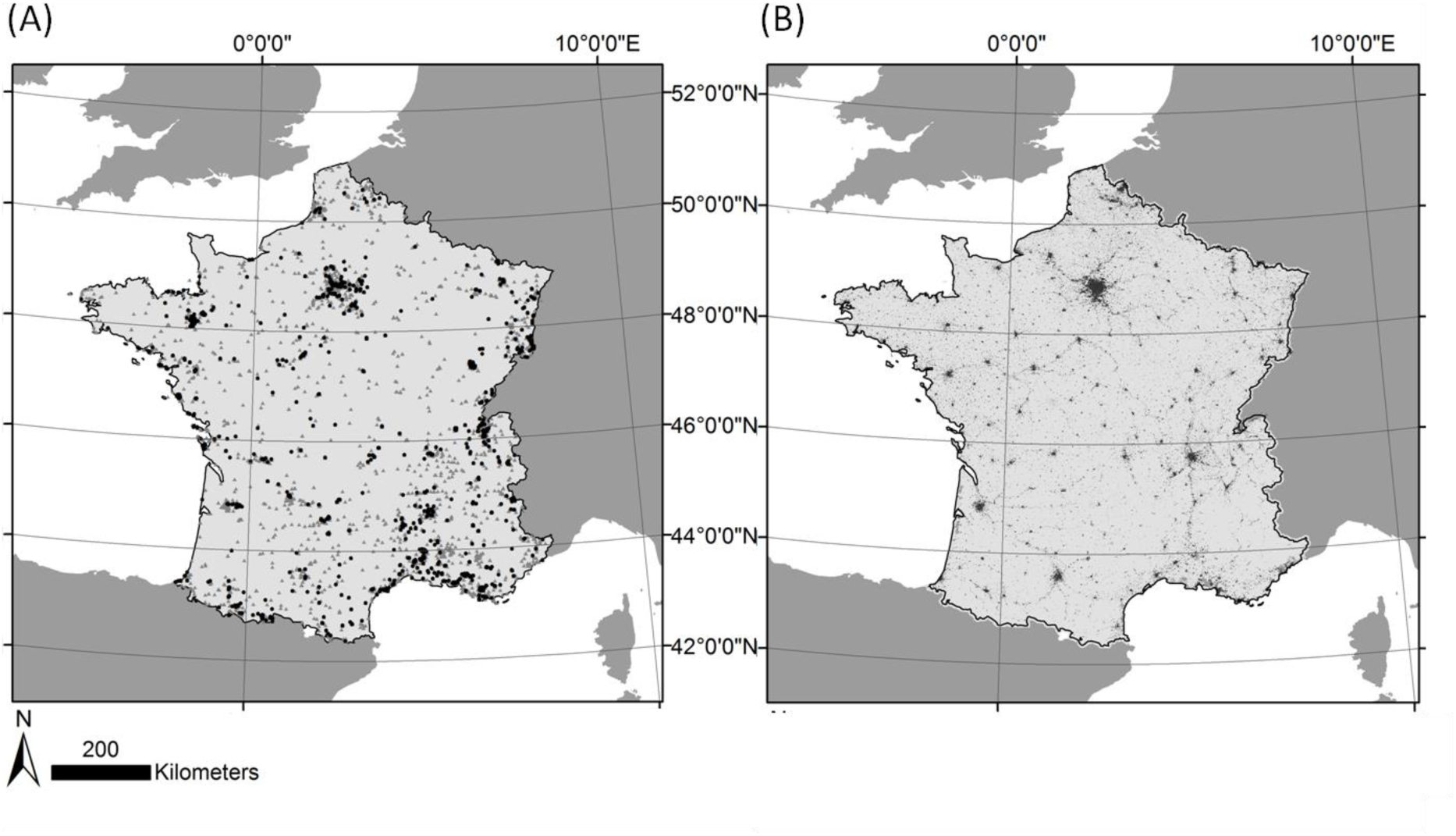
Locations of collections and urbanization in Continental France. In both maps, Continental France is represented in light grey and the other emerged areas are represented in dark grey. (A) Collections with ant are represented by black dots and without by grey triangles. (B) Urbanized areas are represented in black and other land uses in light grey.

### 2. The dataset

Data comes from a citizen science program aiming at monitoring flower visitor across France using a standardized protocol fully described elsewhere (Deguines *et al.* 2012). Briefly, volunteers choose a flowering plant anywhere in France and take pictures of every insect visiting its flowers within a 20-minute period. Then, insects and plant pictures are identified using online computer-aided identification tools (Causse *et al.* 2013) that propose sets of predefined taxa or morphospecies, including one “ant” taxa. The date, time, temperature and precise location of the observations are provided by volunteers when uploading their data on the program website (www.spipoll.org). Each set of identified plant and insects visiting it at a given time and place is referred as a collection. For the purpose of the present study, we used all the collections recorded from 2010 to 2014 that comprised a total of 14,027 collections including 2518 picture of ants (Fig 1. A). They spanned a latitudinal gradient from 42.36° to 51.06°N and covering more than 99 % of France latitudinal gradient. All ant pictures were then identified to the highest taxonomic resolution possible by professional experts. S1 Table in Supporting Information detailed these taxonomic groups, ranging from the family to the species level.

### 3. Urbanization index

Urbanization was characterized by the artificial area category of the first level of Corinne Land Cover 2006 database (raster version 25 m resolution, EEA 2010). In France, urbanization is positively correlated with latitude (EEA 2006). To disentangle the potential urbanization and latitude effects, we followed the methodology proposed by Deguines *et al.* (Deguines *et al.* 2012) and calculated a relative urbanization index. Precisely, for each collection, we first measured the proportion of urban land use within a 1 km radius. Then, for each collection we subtracted to this proportion the mean proportion of urban land use within a 1 km radius of all the collection located within a 100 km radius of the focal collection. This index of relative urbanization is thus positive when the focal collection is more urbanized than the collections that have been sampled regionally (i.e. within a 100 km radius) and negative when the focal collection is less urbanized than the collection sampled regionally. Seven Collections that presented less than 30 other collections in the 1 km radius where excluded from the analysis.

### 4. Statistical Analysis

We analysed the presence/absence of the nine most abundant ant taxonomic groups within the collections using generalised linear models with binomial error distribution. We included as explanatory variables in the model the latitude of the collection, its index of relative urbanization and the interaction between both. To allow for quadratic response to latitude, we also included the squared latitude and its interaction with the index of relative urbanization. All models were simplified to the minimum adequate models based on the Akaike Information Criterion (Johnson and Omland 2004). Significant of parameters was obtained with an Anova type 3 test.

## Results

### 1. Identity of floricolous ants

Flower visiting ants were present in 17.9 % of the 14,027 collections included in our dataset, totalising 2511 ant observations. We specified the identification of 92 % of these observations to a level ranging from subfamily to species (Table 1 and S1).

**Table 1.** Occurrences of the 10 taxa studied.

| Taxa levels |  |  |  |
| --- | --- | --- | --- |
| family | subfamily | Genus | Species |
| Ants<br>(Formicidae,<br>2511) | Dolichoderinae (116) |  |  |
|  | Formicinae (2061) | *Camponotus* (464) | *Camponotus lateralis* (Olivier, 1792)<br>(76) |
|  |  |  | *Camponotus piceus* (Leach, 1825)<br>(120) |
|  |  | *Lasius* (781) | *Lasius emarginatus* (Olivier, 1792)<br>(116) |
|  |  | *Serviformica* (697) |  |
|  | Myrmicinae (174) |  |  |

#### Authority are given for species level

This revealed that floricolous ants were not evenly distributed among subfamilies: 81 % belonged to Formicinae, 7 % to Myrmicinae and 5 % to Dolichoderinae (Fig 2 (A)). The remaining 8 % of ant picture were not good enough to attribute them to a subfamily. Within Formicinae, three genera were highly dominant: *Lasius, Serviformica* (subgenus of *Formica*) and *Camponotus* that represented respectively 31 %, 27 % and 18 % of all ant observations (Fig 2 (A)). Within these genera, we were further able to identify three species: *Lasius emarginatus, Camponotus piceus* and *Camponotus lateralis* that represented respectively 4.6 %, 4.7 % and 3.0 % of all identified ants.

**Fig 2.**
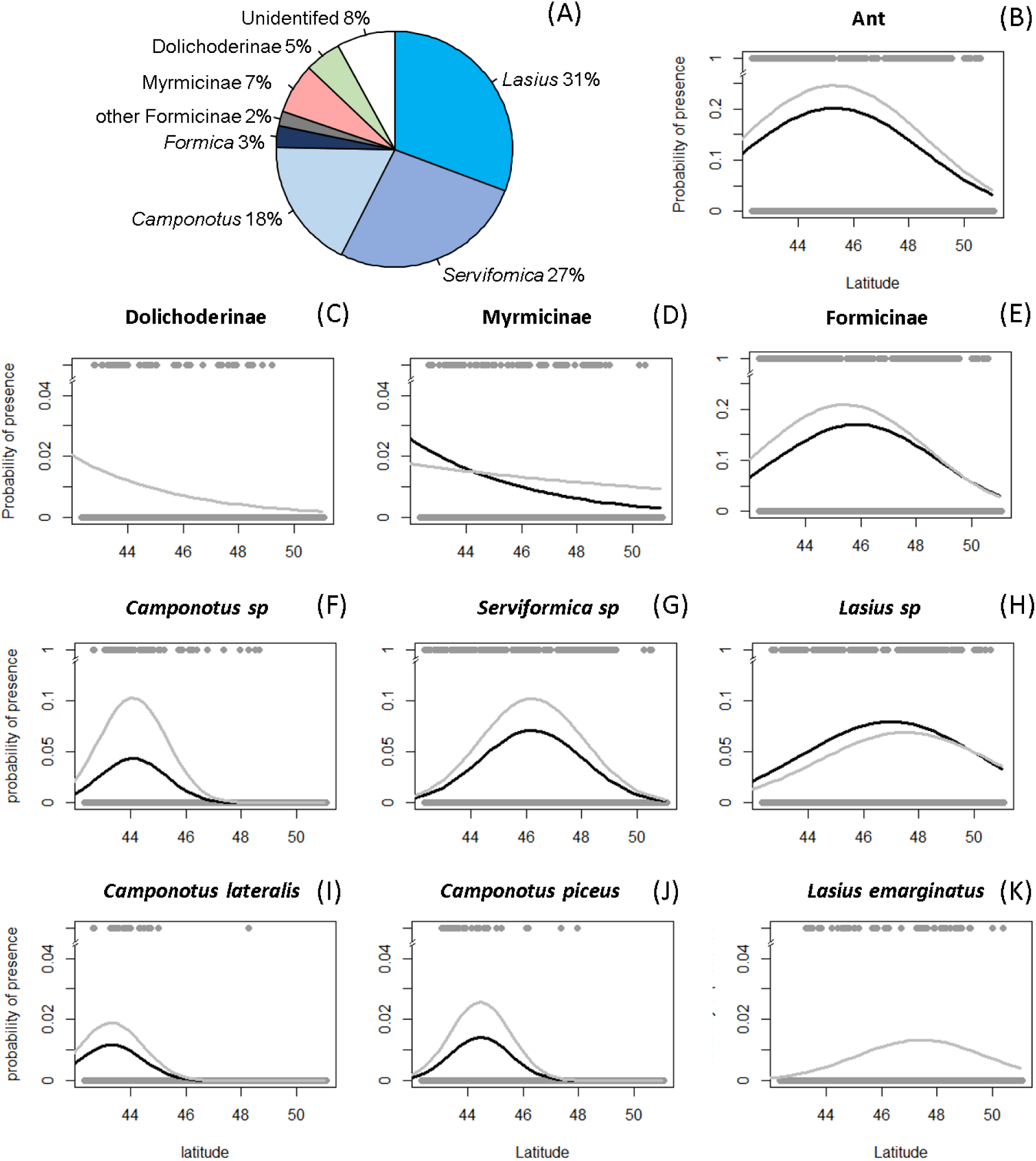
Pie chart of the relative presence and probability of presence along latitudinal and urbanization gradients of the nine taxa studied. (B to K) Lines represent prediction from generalized linear models similar to the ones presented in the methods except that the relative urbanization effect was treated as a factor with two levels: low (grey) and high (black), each corresponding to half of the collections with respectively the lowest and highest values of relative urbanization.

### 2. Responses to latitudinal and urbanization gradients

When analysing the distribution of ant observations across France, we found a significant quadratic effect of the latitude indicating a hump-shaped response of ant presence with an optimum around 45-46 °N (Fig 2 (B) and Table 2). We further found a significant interaction between the latitude and the index of relative urbanization indicating that urbanization affected ant presence differently along the latitudinal gradient, with a negative effect at lower latitude (Fig 2 (B) and Table 2).

**Table 2.**
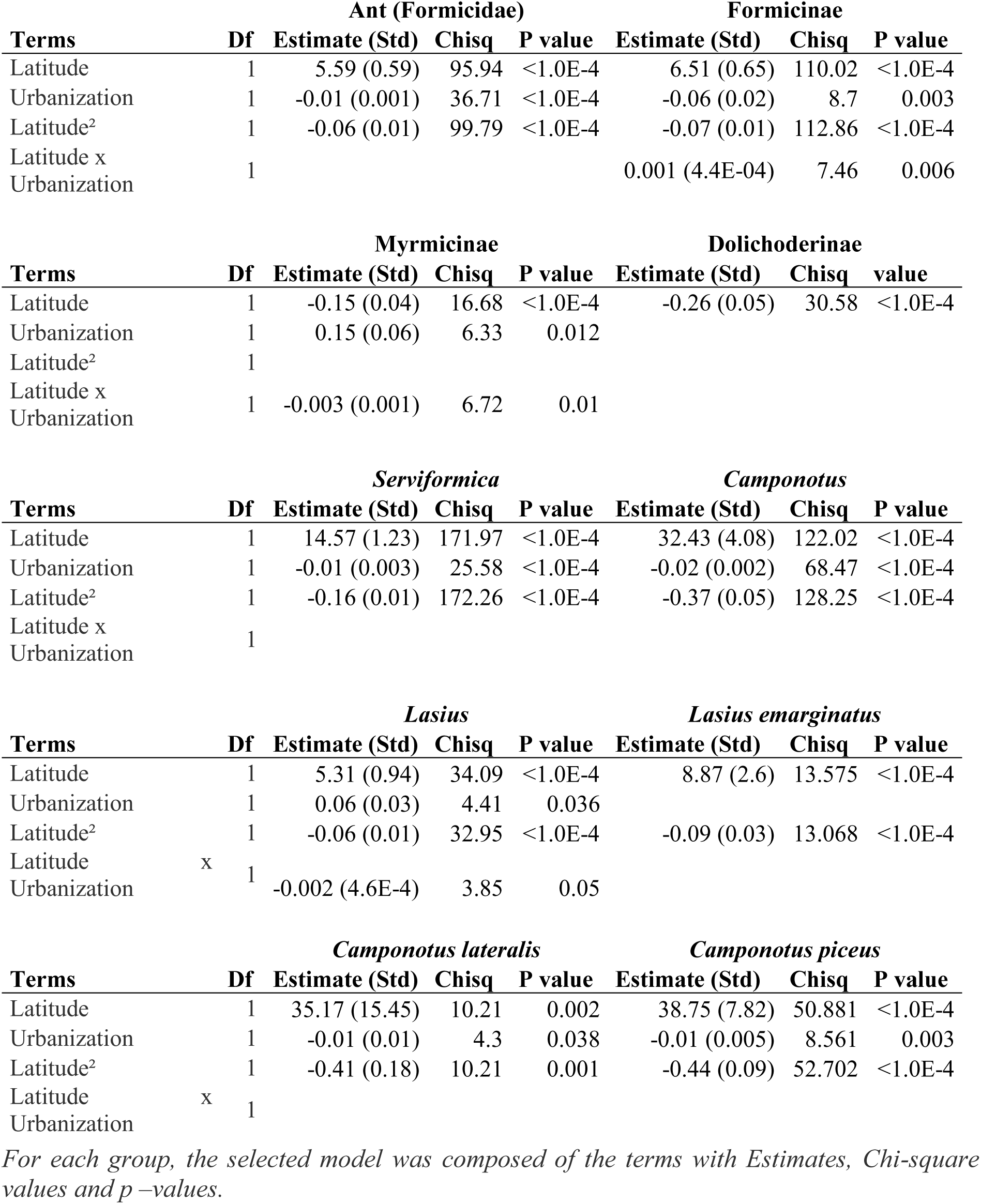
Minimum adequate models for each of the nine ant groups.

The analysis of the presence of the three ant subfamilies highlighted striking differences among them. First, Formicinae subfamily exhibited a pattern similar to the one observed for all the ants, with a significant quadratic effect of the latitude with an optimum around 45-46 °N, and a significant interaction between the latitude and the index of relative urbanization showing a negative impact at low latitude (Fig 2 (E) and Table 2). Second, Myrmicinae subfamily exhibited a significant interaction between the latitude and the index of relative urbanization highlighting a positive impact at low latitude and a negative impact at high latitude (Fig 2 (D) and Table 2). Finally, for Dolichoderinae subfamily, we found a significant effect of the latitude (Table 2) revealing a steady decrease in the presence of this subfamily with increasing latitude (Fig 2 (C)).

Scaling down our analysis at the genus level, we found a significant quadratic effect of latitude for the three main genera belonging to Formicinae subfamily (Table 1). Interestingly, the optimum was different for each genus. While *Camponotus* appeared to be the most southern genus with a presence peak around 44 °N, the optimum for *Serviformica* was around 46 °N and *Lasius* appeared to be the northernmost genus with its optimum located between 46 and 48 °N (Fig 2 F, 2 G and 2 H, respectively). The presence of these three genera was also significantly affected by the level of relative urbanization, but slightly differently. For *Serviformica* and *Camponotus* genus, we found a significant negative effect of the relative urbanization (Fig 2 (F) and 2 (G) and Table 2). For *Lasius*, we found a significant interaction between the relative urbanization and the latitude indicating that the effect of urbanization was dependent on the latitude, with a positive effect for intermediate latitudes (between 45-47 °N) (Fig 2 (H) and Table 2).

Scaling further down our analysis to the three species we could identify and with sufficient data (S1 appendix), we found a significant quadratic effect of latitude for each one. As for the three genera analysed above, we found that the optimum varied among species: *Camponotus lateralis* appeared to have the southernmost distribution, peaking around 43.5 °N (Fig 2 (I)), followed by *Camponotus piceus* that peaked between 44 and 44.5 °N (Fig 2 (J)). *Lasius emarginatus* had the most northern and spread out distribution that peaked around 47.5 °N (Fig 2 (K)). While the level of relative urbanization did not affect *Lasius emarginatus* (Fig 2 (K)), we found a significant negative effect of the relative urbanization for *Camponotus lateralis* (Fig 2 (F) and Table 2) and *Camponotus piceus* (Fig 2 (J) and Table 2).

## Discussion

### 1. Dominance of Formicinae on flowers

At the subfamily level, almost 80 % of the observed floricolous ants belonged to Formicinae subfamily. This result is consistent with Rico-Gray and Oliveira (Rico-Gray and Oliveira 2007) that considered Formicinae as one of the most floricolous subfamily which could be explained by the liquid-feeding (flower nectar or honeydew from aphids) behaviour of most species. Surprisingly, ants from the subfamily Dolichoderinae, which is also considered as highly floricolous (Rico-Gray and Oliveira 2007) were observed in 5 % of the collections only. This overall low occurrence could be explained by the mainly southern distribution observed; this pattern is consistent with what is known of the distribution of this subfamily (Blatrix *et al.* 2013). Turning to Myrmicinae, their low occurrence in our dataset is consistent with their more predators and less floricolous habits than the two previous subfamilies (Rico-Gray and Oliveira 2007). Finally, the negligible occurrences of the last three subfamilies present in France, the Ponerinae, (less than 0.1 %), none of the Leptanillinae nor the Proceratiinae is again consistent with their almost strict edaphic behaviour in Continental France (Bernard 1968, Blatrix *et al.* 2013).

### 2. Match and mismatch with known species latitudinal distribution

Most of the knowledge about ant distribution is at the species level. Two of the three ant species with sufficient data to analyse their latitudinal distribution on flower showed a good match between the distribution we observed and their known latitudinal species distribution. Indeed, the narrow and southern range we observed for *C. lateralis* is consistent with Blatrix *et al.* (2013) that consider this species as absent or very rare above 46 °N. Similarly, the wide latitudinal range of *L. emarginatus* we observed on flowers is consistent with its known presence all over Continental France (Blatrix *et al.* 2013).

Interestingly presence on flowers of *C. piceus*, the last ant species with enough data to analyse its latitudinal distribution, was observed on a much narrower latitudinal range than its known distribution (between 43-50 °N, Blatrix *et al.* 2013). For ants, inter-specific competition is considered as the major driver of community composition (reviewed by Cerdá *et al.* 2013) and especially competition for food resources (Albrecht and Gotelli 2001, Blüthgen and Fiedler 2004, Lester *et al.* 2010, Vepsäläinen and Pisarski 1982). Vepsäläinen and Pisarski (1982) divided species in three levels of hierarchy based on their respective dominance in communities: (1) dominants that compete and exclude other species from territory and food resources (2) subdominants that aggressively defend or try to take over food resources and (3) subordinates that defend only their nest. However, most ant assemblage lack of species from the dominant category and some subdominant species clearly act as dominant (Cerdá *et al.* 2013). This pattern has been observed for *L. emarginatus* in the habitats of Crimea Mountains where its colonies were the largest (around 30 000 workers, Stukalyuk and Radchenko 2011) and dominated other species such as *C. piceus.* In our case, dominant and almost strict liquid feeding species such as *L. emarginatus* (Blatrix *et al.* 2013) could strongly compete for the food resources produced by the plants (directly such as nectar or indirectly such as aphids honeydew) and outcompete less dominant and more diet generalist species such as *C*. piceus (Blatrix *et al.* 2013).

### 3. Segregate and nested responses to latitude at different taxonomic levels

Overall, floricolous ants showed a quadratic response to latitude, with maximum occurrence at intermediate latitudes (around 45-46° N). Such pattern echoed the one previously described on the species richness of coprophageous beetles studied along the same latitudinal range (Lobo *et al.* 2002). According to Lobo *et al.* (2002), this pattern could be explained by maximum annual and mean annual temperatures which are the most in?uential spatially structured climate variables along latitudinal gradient of continental France.

We highlighted that the presence of the three main subfamilies of floricolous ants exhibited different latitudinal optimums, with Myrmicinae and Dolichoderinae being restricted to lower latitudes whereas Formicinae were more present at medium and high latitudes. Strikingly such a segregated response of ant to latitude was also observed for the lower taxonomical levels we studied, with the presence of the 3 genera of Formicinae showing segregated optimum along the latitudinal gradient (from south to north France: *Camponotus, Serviformica* and *Lasius*) and the two species of *Camponotus* showing the same pattern (from south to north: *C. lateralis* and *C. piceus*). Such pattern could result from niche partitioning among ant taxa to minimise competition for resources (reviewed by Schoener 1974), a phenomenon that has been shown to be related to the distribution of ants along large scale environmental gradient (Machac *et al.* 2011). However, we also observed that the latitudinal segregation was taxonomically aggregated which could be the sign of shared ancestral ecological characteristics. However, more data on various ant species are needed to properly test for a phylogenetic signal in the latitudinal optimum of ants.

### 4. Heterogeneous responses to Urbanization and interaction with latitudinal gradient

Most ant taxa analysed here showed a negative response to urbanization gradient. This is in accordance with previous studies highlighting a negative impact of urbanization on ant abundance (Buczkowski and Richmond 2012) or richness (McKinney 2008, Vepsäläinen *et al.* 2008). Such a negative effect has also been found for other floricolous arthropods, such as Lepidopterans, Coleopterans and Dipterans (Deguines *et al.* 2012), but also for ground beetles (Vergnes *et al.* 2014), suggesting that the effect of urbanization on arthropods is general.

However, the lower presence of ant visiting flowers in urbanized areas could also be link to a decrease in the “attractivity” of flowers as resources. Urbanization strongly modifies the environment (i.e. soil and air properties, carbon cycle and management) (McDonnell and Hahs 2008). These modifications could have altered the properties of nectar (attractiveness, quality or quantity) (Clark *et al.* 2007, Meindl and Ashman 2013). Moreover, cities are highly heterotrophic systems that produces a lot of resources for invertebrates (McIntyre 2000) such as wastes that are known to be highly attractive for ants (Youngsteadt *et al.* 2014). Ant might thus neglect floral resources in urban environment, to the profit of other urban specific resources. Finally, exclusive competition by strict liquid eating and dominant ants such as *L*. *emarginatus* could reinforce such a switch in food resource for omnivorous and less dominant species such as *C. piceus* or *C. lateralis* (Blatrix *et al.* 2013, Cerdá *et al.* 2013).

As expected, we observed a contrasted response to urbanization along the latitudinal gradient with a tendency for a stronger negative effect at lower latitudes. The increase in temperature and decrease of soil moisture along urbanization gradient are considered as general trends (McDonnell and Hahs 2008). However, in regions with a Mediterranean climate such as the lowest latitudinal parts of France, an increase of soil moisture and a decrease of soil temperature along urbanization gradient is frequently observed, mainly due to run-off on impervious surfaces (Holway and Suarez 2006). Those colder and moister conditions are less favorable to most ants, as observed at large scale across Europe (Kumschick *et al.* 2009).

## Conclusion

Our analysis highlights that both latitude and urbanization affect the distribution of ant-flower interactions and that the coupling of a large dataset collected by citizen scientists with appropriate data validation by experts is an efficient and promising approach. Our results suggested a strong impact of both evolutionary history and competition in shaping the distribution of ant plant interaction across France.

## Declarations

We wish to thanks the hundreds of participants of the Spipoll program who collected the data and Mathieu de Flores (OPIE) for coordinating this program.

## Conflict of interest disclosure

The authors of this article declare that they have no financial conflict of interest with the content of this article.

## Supporting information

Additional Supporting Information may be found in the online version of this article: **S1 Table**. Details of taxonomic levels between studied groups.

